# Soft Windowing Application to Improve Analysis of High-throughput Phenotyping Data

**DOI:** 10.1101/656678

**Authors:** Hamed Haselimashhadi, Mason C. Jeremy, Violeta Munoz-Fuentes, Federico López-Gómez, Kolawole Babalola, Elif F. Acar, Vivek Kumar, Jacqui White, Ann M. Flenniken, Ruairidh King, Ewan Straiton, John Richard Seavitt, Angelina Gaspero, Arturo Garza, Audrey E. Christianson, Chih-Wei Hsu, Corey L. Reynolds, Denise G. Lanza, Isabel Lorenzo, Jennie R. Green, Juan J. Gallegos, Ritu Bohat, Rodney C. Samaco, Surabi Veeraragavan, Jong Kyoung Kim, Gregor Miller, Helmut Fuchs, Lillian Garrett, Lore Becker, Yeon Kyung Kang, David Clary, Soo Young Cho, Masaru Tamura, Nobuhiko Tanaka, Kyung Dong Soo, Alexandr Bezginov, Ghina Bou About, Marie-France Champy, Laurent Vasseur, Sophie Leblanc, Hamid Meziane, Mohammed Selloum, Patrick T. Reilly, Nadine Spielmann, Holger Maier, Valerie Gailus-Durner, Tania Sorg, Masuya Hiroshi, Obata Yuichi, Jason D. Heaney, Mary E Dickinson, Wurst Wolfgang, Glauco P. Tocchini-Valentini, Kevin C. Kent Lloyd, Colin McKerlie, Je Kyung Seong, Herault Yann, Martin Hrabé de Angelis, Steve D.M. Brown, Damian Smedley, Paul Flicek, Ann-Marie Mallon, Helen Parkinson, Terrence F. Meehan

## Abstract

**Motivation:** High-throughput phenomic projects generate complex data from small treatment and large control groups that increase the power of the analyses but introduce variation over time. A method is needed to utlize a set of temporally local controls that maximises analytic power while minimising noise from unspecified environmental factors.

**Results:** Here we introduce “soft windowing”, a methodological approach that selects a window of time that includes the most appropriate controls for analysis. Using phenotype data from the International Mouse Phenotyping Consortium (IMPC), adaptive windows were applied such that control data collected proximally to mutants were assigned the maximal weight, while data collected earlier or later had less weight. We applied this method to IMPC data and compared the results with those obtained from a standard non-windowed approach. Validation was performed using a resampling approach in which we demonstrate a 10% reduction of false positives from 2.5 million analyses. We applied the method to our production analysis pipeline that establishes genotype-phenotype associations by comparing mutant versus control data. We report an increase of 30% in significant p-values, as well as linkage to 106 versus 99 disease models via phenotype overlap with the soft windowed and non-windowed approaches, respectively, from a set of 2,082 mutant mouse lines. Our method is generalisable and can benefit large-scale human phenomic projects such as the UK Biobank and the All of Us resources.

**Availability and Implementation:** The method is freely available in the R package SmoothWin, available on CRAN http://CRAN.R-project.org/package=SmoothWin.

## Introduction

High-throughput, large scale phenotyping studies evaluate variables of an organism’s biological systems to examine the contribution of genetic and environmental factors to phenotypes. Standardised phenotyping screens that cover a wide range of biological systems have made useful insights for identifying new genetic contributors to robust phenotypes as compared to more focused studies that often target well-characterised genes with varying reproducibility ^1–5^. Leveraging economies of scale and using standardised procedures, high-throughput phenotyping screens addresses these challenges and have been applied in biological screening of chemical compound libraries, agricultural evaluation of crop plants, genome-wide CRISPR-based mutagenic cell line screens and multi-centre phenotypic screening of mutated model organisms ^6–13^. The continuous generation of large volumes of data introduces new challenges affecting automated approaches to statistical analysis that have to scale with increasing data and address the underlying complexity inherent in large projects ^14–17^.

The International Mouse Phenotyping Consortium (IMPC) is a G7 recognised global research infrastructure dedicated to generating and characterising a knockout mouse line for every protein-coding gene ^18–20^. Currently, the IMPC has phenotyped over 148,000 knockouts and 43,000 control mice (data release 9.2, January 2019) across 11 research centres in 9 countries. These centres adhere to a set of standardised phenotype assays defined in the International Mouse Phenotyping Resource of Standardised Screens (IMPReSS), and designed to measure over 200 parameters on each mouse. As part of these standardised operating procedures, critical factors that can impact data collection, such as reagent type or equipment, are reported as required metadata. Phenotype data is then centrally collected and quality controlled by trained professionals before being released for analysis. All phenotype data is processed by the statistical analysis package PhenStat—a freely-available R package that provides a variety of statistical methods for the identification of genotype to phenotype associations by comparing mutant to control data that have the same critical attributes ^17^. For quantitative data, linear mixed models are typically employed with several factors modelled in including sex, sex-genotype interaction, body weight, and batch (i.e., phenotype measures collected on the same day). Mutant mouse lines found to have a significant deviation in phenotype measurements are assigned a phenotype term from the Mammalian Phenotype Ontology ^21^. These associations, as well as the raw data, are disseminated via the web portal (https://www.mousephenotype.org) using application programming interfaces (APIs) and data downloads.

A challenge with high-throughput phenotyping efforts is the small sample size for the experimental group (i.e., the knockout mice) that is produced to maximise the use of finite resources, considering biological relevance and power analysis ^22^. The IMPC centres are encouraged to measure these knockout mice in two or more batches, as this improves the false discovery rate by modelling in the random effect of day-to-day variation ^23^. In contrast, large control sample sizes accumulate as they provide a strong internal control of the pipeline and typically generated with every experimental batch. Such large control groups represent a unique dataset that increase the power of the subsequent analyses and allow the construction of a robust baseline^19^. However, this can lead to the accumulation of heterogeneities including seasonal effects, changes in personnel, and unknown time-dependent environmental factors ^23^.

A simple approach to cope with heterogeneity in the data is to set explicit time boundaries (e.g., one year) before and after experimental collection dates. This “hard windowing” approach will capture different time-frames depending on how much time elapses between the first and last batch of experimental data measured. This approach is unsatisfactory for IMPC data as some mutant lines had enough experimental mice to measure in one batch, while others needed multiple batches over 18 months due to breeding difficulties or other factors. This variation in time-frames can lead to a widely different number of controls being applied to an analysis, making it challenging to explore correlations between mutant lines. Thus, more tunable approaches were needed.

In this study, we address the complexity of the data collected over time by proposing a novel windowing strategy that we call “soft windowing”. This approach utilises a weighting function to assign flexible weights, ranging from 0 to 1, to the control data points. Controls that are collected on or near the date of mutants are assigned the maximal weights, whereas controls at earlier or later dates are assigned less weight. In contrast to the hard windowing, the weighting function in this soft windowing allows for different shapes and bandwidths by alternating the tuning parameters. In addition, we demonstrate how to tune parameters and demonstrate the implementation of the soft windowing on IMPC data.

### System and methods

In high-throughput projects, such as the IMPC, the model parameters may not stay constant over time that can lead to misleading inferences. For example, Figure 1 illustrates changes to the control group trend and/or variation over time for the *Forelimb grip strength normalised against body weight* and *Mean cell volume*. One approach widely used in signal processing ^24–27^ is to define a windowing function that includes the appropriate number of data points to capture the effect of interest while minimising the noise. This is defined by

**Fig 1:**
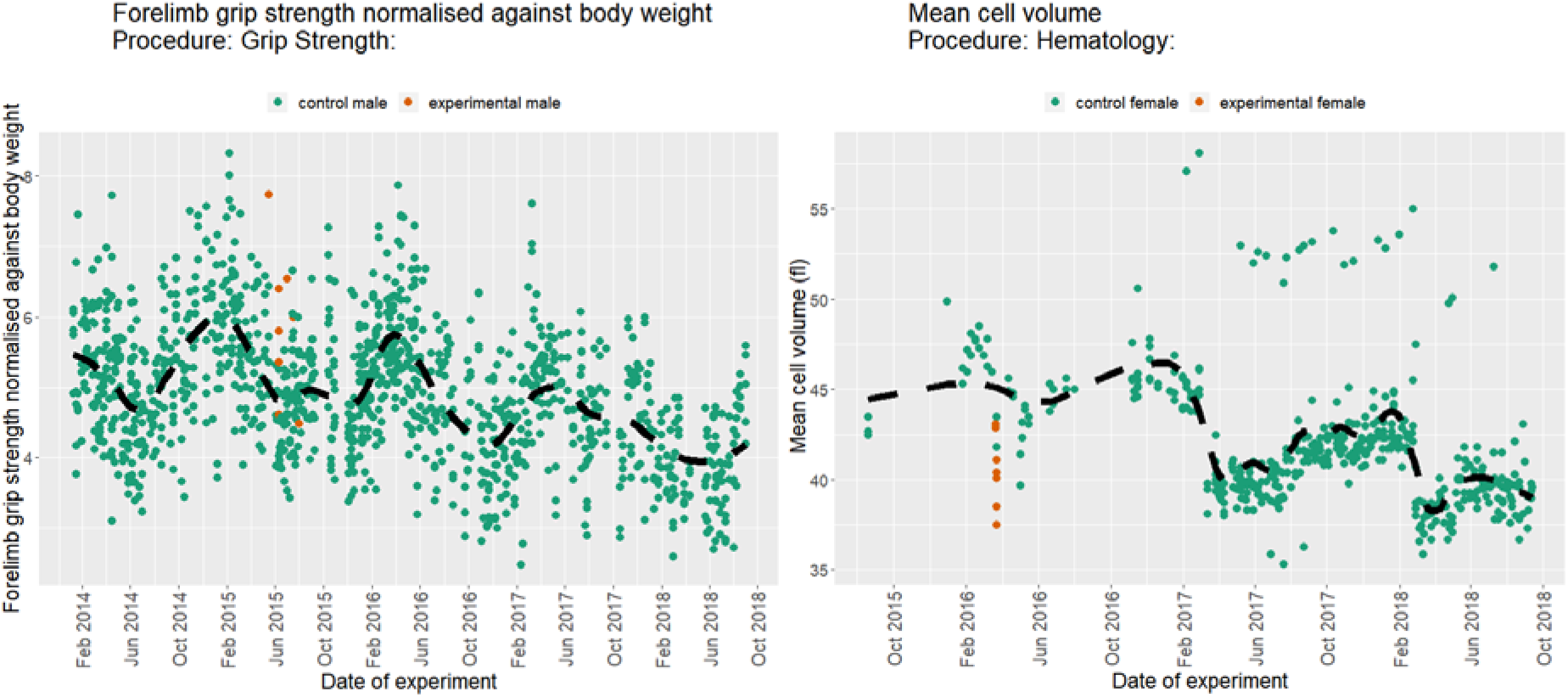
Examples of longitudinal data from the IMPC selected for high variance in control population. Scatter plot of the *Forelimb grip strength normalised against body weight* (left) and *Mean cell volume* (right) from the IMPC Grip Strength and Haematology procedures, respectively. The dashed black line represents the overall trend of the controls (dark blue). Mutant mice are in orange.

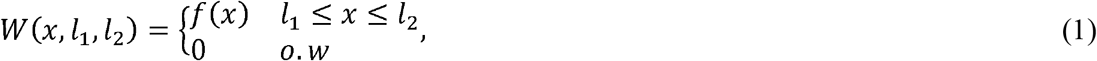

where setting *f*(*x*) to a constant, e.g., *f*(*x*) = 1, leads to hard windowing, while setting it to a smooth function results in the soft windowing. The same approach can be generalised to multiple signals ^28–30^ or applied as a rolling window ^31^ in the presence of exogenous variables to account for time dependency in the regression coefficients ^32^. Alternatively, we propose a soft windowing approach for the regression methods by defining a weighting function that applies less weight to the residuals outside the window of interest. This leads to distinct advantages over the hard windowing. First, the entire dataset is included in the analysis in contrast to the limited data points in the hard windowing. Second, the windowing and the parameter estimation are coupled, which is a direct result of using the Weighted Least Squares (WLS). Critically, by bounding the controls in a window, we freeze the analysis and abrogate the need for further analysis assuming no new experimental data is generated within the time window.

## Algorithm

Our novel windowing strategy explicitly defines the weighting function and proposes a simple but effective set of criteria to estimate the minimal window for the noise-power trade off.

### Weight generating function

Let *t* = (*t*_1_, *t*_2_,…, *t*_*n*_) represent a set of *n* continuous time units, *m* = (*m*_1_, *m*_2_,…, *m*_*p*_) the time units when the treatments are measured (peaks in the windows), *l* = {(*l*_1*L*_,*l*_1*R*_), (*l*_2*L*_,*l*_2*R*_), …, (*l*_*pL*_,*l*_*pR*_)} a set of *p* non-negative left and right *bandwidths* and *k* = {(*k*_1*L*_, *k*_1*R*_), (*k*_2*L*_,*k*_2*R*_), …, (*k*_*pL*_, *k*_*pR*_)} a set of *p* positive left and right *shape* parameters. We impose the continuity on the time to simplify the definition of a continuous function over the time units, e.g., by converting dates to UNIX timestamps. Furthermore, we introduce a peak generating function (PGF) of the form of *c*_*i*_ = *F*(*t*;*m*_*i*_ − *l*_*iL*_,*k*_*iL*_)(1 − *F*(*t*;*m*_*i*_ + *l*_*iR*_,*k*_*iR*_)), *i* = 1,2, …,*p* where *F*(*x*; *μ,σ*) = *Pr*_*X*_(*X* ≤ *x*|*μ,σ*) is selected from the family of cumulative distribution functions (cdf) with location *μ* and scale *σ*. In this study, we select *F* from the family of continuous and symmetric distributions (such as the Logistic, Gaussian, Cauchy and Laplace distributions). Then, we propose a weight generating function (WGF) of the form of

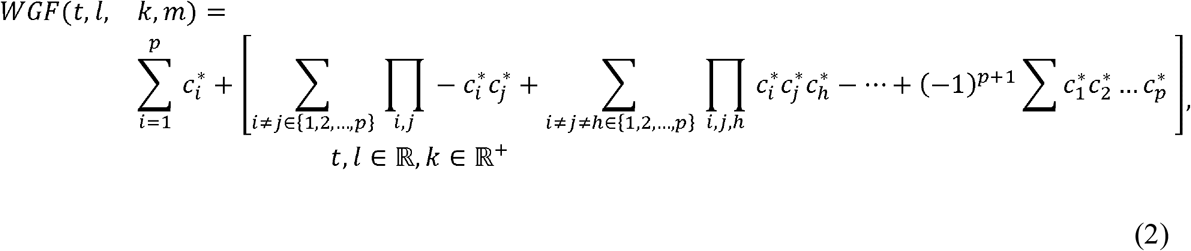

where 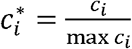 denotes the normalised peak generating function. The first term on the right hand side of Eq. 2 produces the individual windows and the second term accounts for merging the intersections amongst the windows. Figure 2 shows the symmetric weight generating function (SWGF), that is *l*_*iR*_ = *l*_*iL*_ and *k*_*iR*_ = *k*_*iL*_,*i* = 1,2, …,*p*, for the different values of *k* ∈ [0.2,50] coloured from blue *(k* = 50) to red *(k* = 0.2) and for the different values of *l* = 5,10,15. The vertical black dashed lines show the hard window corresponding to the value of *l*. From this plot, the function is capable of generating a range of windows from hard (blue) to smooth (red). Further, the weights lay in the (0,1] interval for all values of time; however, they may not cover the entire (0,1] spectrum in a bounded time domain. Then, the weights are normalised to be ranged in (0,1] before inserting into the WGF as shown by 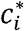 in Eq 2. Figure 3 shows the merge capability of the SWGF for the logistic *F* with *m* = 15,35 and different values of *k* = 0.5, 1.5,3 and *l* = 6,8,10,12. From this figure, the function is capable of producing a range of flexible multimodal windows (top) as well as aggregated windows (bottom) if |*m*_1_+*l*| > |*m*_2_ − *l*| for all *m*_1_ < *m*_2_, *l* ∈ ℝ. In all cases, the weights lay in the (0,1] interval.

### Windowing regression

Let *y* = *xβ* + *e* denote a linear model, with *y, x, β* and *e* representing response, covariates, unknown parameters and independent random noise, *e* ~ *N*(0, *σ*^2^ < ∞) respectively. Imposing the weights in Eq. 2 on the residuals leads to the following weighted least square (WLS)

**Figure 2:**
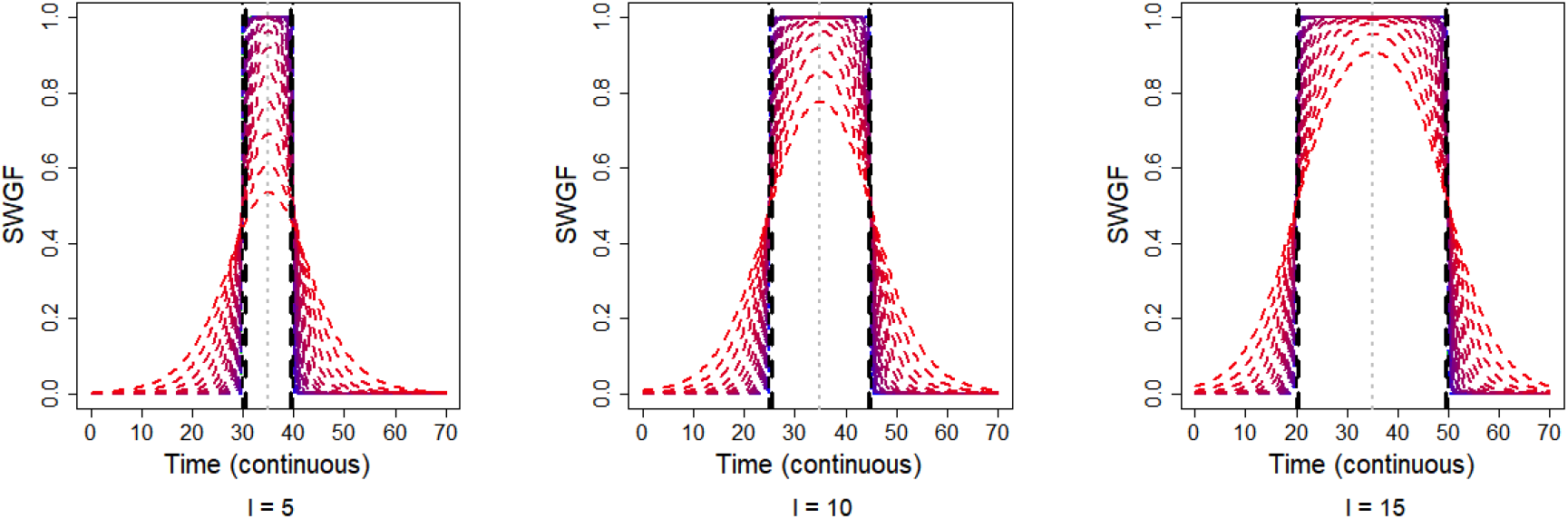
Behaviour of the Symmetric Weight Generating Function (SWGF) for a spectrum of values for the shape parameter, *k*, ranging from *k* = 50 (blue) to *k* = 0.2 (red), in intervals of *t* = 1,2,…,70, and for the different values of the bandwith *l* = 5,10,15 (left to right). The black dashed lines show the hard windows corresponding to *l*. The gray dotted lines show the window peaks. These plots show the capability of the WGF to generate different forms of the window.

**Fig 3:**
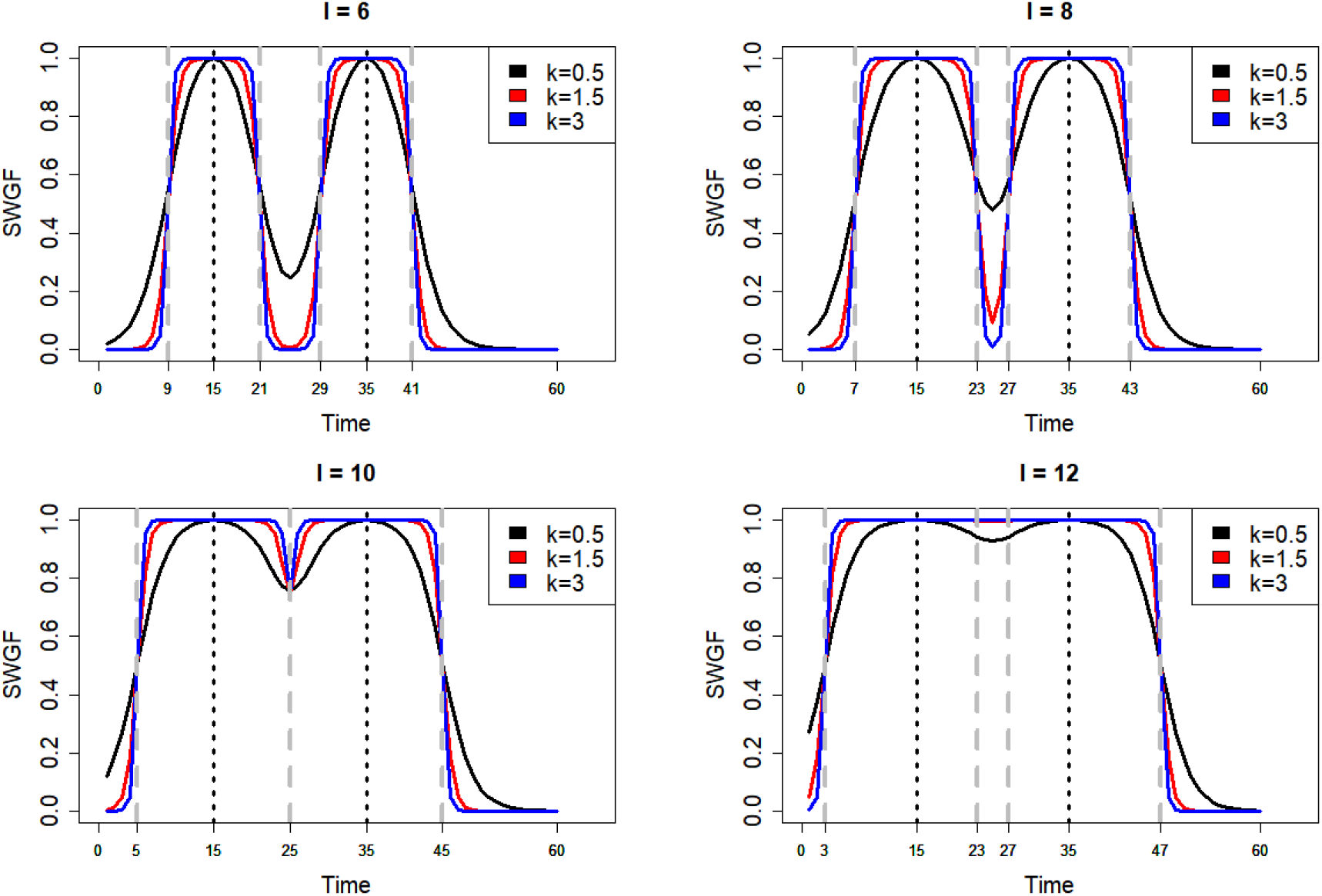
Merging behaviour of the SWGF for different values of the shape parameter *k* = 0.5,1.5,3 and the bandwidth *l* = 6,8,10,12 on a sequence of time points *t* = 1,2,…, 60. The vertical dashed gray lines show the corresponding hard windows to *l*. This plot shows the capability of SWGF to generate multimodal windows as well as merging individual windows.

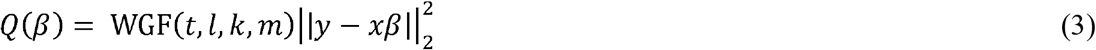

Where |ǀ⋅ǀ|_2_denotes the second norm of a vector. Minimising *Q(β)* with respect to *β* leads to 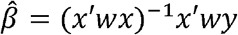, where *w* is a diagonal matrix of weights from WGF and (′) denotes the transpose of a matrix. Weighted linear regression (WLR), in the context of this study, is equivalent to imposing less weight on the off modal time points with respect to *m.* We illustrate this in Figure 4, where 60 observations are simulated from the following model,

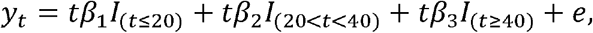

with *t* = 1,2,…,60, *β*_1_ = 0, *β*_2_ = 1, 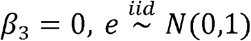 and *I* is the indicator function,

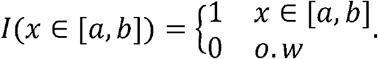

In other words, the model is piecewise linear and only significant in the *t* ∈ (20,40) interval. Figure 4 (left) shows the global estimation of the linear regression from the entire data (dotted black line) and the WLR by WGF (*t*,9,5,30) (dashed blue line) as well as weights from the WGF on the right. This plot shows that the non-weighted linear regression leads to a horizontal line, where no significant gradient is detected, whereas the WLR tends to model the significant section of the data that leads to fitting the true line. Figure 4 compares the effect of windowing vs. considering the entire dataset, showing the different conclusions.

**Fig 4:**
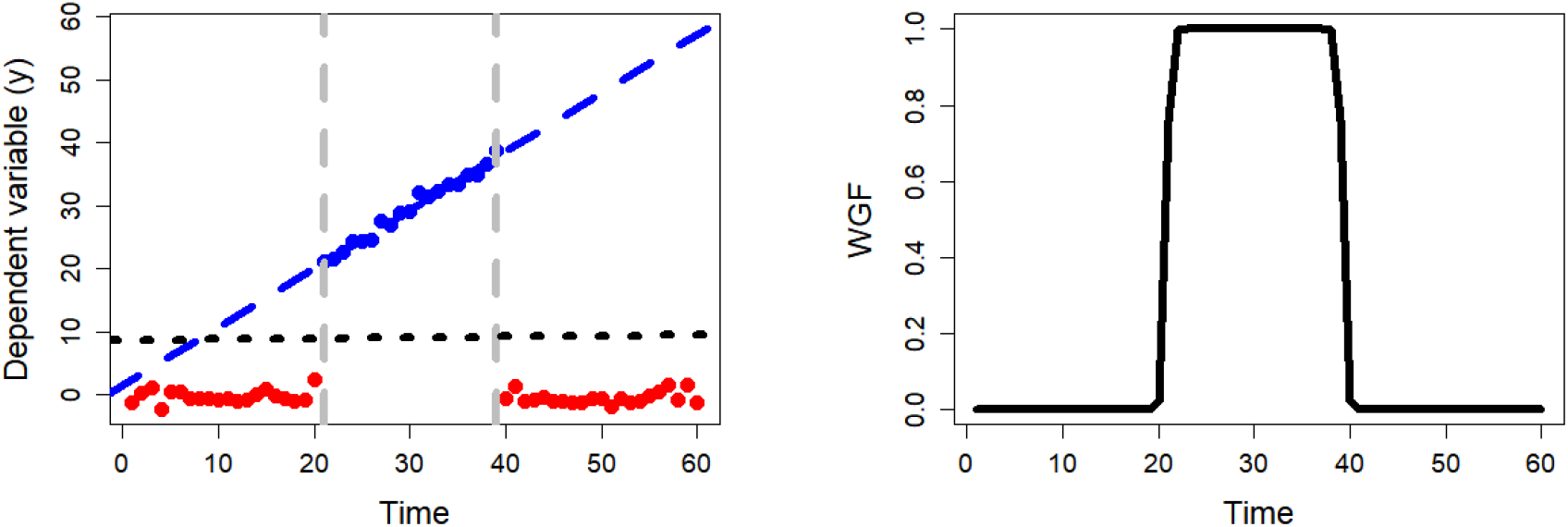
(Left) Comparison between the inferences from the windowed linear regression on the simulated data (blue dashed line) and without windowing (dotted black line). (Right) The corresponding weights from WGF centred on m = 30. With windowing, we attempt to model the effective section of the data (blue dots).

### Selection of the tuning parameters

Selection of the tuning parameters *k* and *l* to define the soft window have a strong impact on the final estimations and consequently on the inferences that are made from the statistical results. Indeed, a wide or over-smooth window can lead to the inclusion of too much noise, whereas a small window can result in low power in the analysis. An additional challenge is the direct linear correlation between increasing the number of peaks, *m*, and to the total number of the parameters for the windows (*l,k*) that results in significant growth in the computational complexity of the final fitting. This is due to tuning the window in the general form of WLS in Eq. 3 requires 2*p* dimensions in space to search for the optimal *l* and *k*. To cope with this complexity, we propose to fix *l* and *k* so all windows are symmetric and have the same shape and bandwidth. We then select the tuning parameters by searching the space on the grid of *l,k* values and look for the most significant change in mean and/or variation of the residuals/predictions. The grid is searched by generating a series of scores from applying t-test (to detect changes in mean) and F-test (to detect change in variation) to the consecutive residuals/predictions at each step of expanding (*l* → *l* + 1) and/or reshaping (*k* → *k* + 1) the windws. This technique is based on the assumption that the mean and the variation of the residuals/predictions remain unchanged in different time periods ^33^.

To gain the necessary power in the analysis, we apply the statistical tests to the values of *l* that correspond to a minimum *T* observations in the windows. Then one can define the quantity of *T*(*l*) that is the total number of observations that is included in the hard window corresponding to *l*. We should stress that the definition of *T*(*l*) in the soft windowing can be challenging because the WGF assigns weights to the entire dataset in the final fitting. To address this complexity, we propose the Sum of Weights Score by 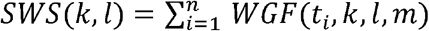, that is the summation of weights from WGF for specific *l* and *k*. Note that *SWS*(*l,k*) ≥ *T*(*l*) with the equality for sufficiently large *k*. Because *l* is generally unknown, a value of *T*(*l*) = *T* independent of *l* needs to be decided before the analysis. Our experiments, inspired by the z-test minimal sample size (*n* > 30), show that setting *SWS* ≥*T* with

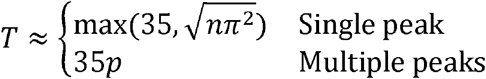

provides sufficient statistical power and precision for the analysis of each sex-parameter in IMPC.

Once the bandwidth,*l*, is selected, the shape parameter, *k*, can be optimised on a grid of values similar to *l*.

This algorithm is implemented for a broad range of models in the R package SmoothWin that is available from https://cran.r-project.org/package=smoothWin. The main function of the package, *SmoothWin*(…), allows an initial model for the input and, given a range of values for the bandwidth and shape, it performs soft windowing on the input model. Furthermore, it allows plotting of the results for diagnostics and further inspections. One also can generate the weights from SWGF using the *expWeigh*(…) function.

## Implementation

### Validation using a resampling approach

To assess the performance of the soft windowing method, we implemented a resampling approach to construct a sample of *artificial mutants* from the IMPC control data by relabelling some control data as mutant. We then examined the difference in the number of false positives that were detected by the standard (non-windowed) analysis versus the soft windowed approach.

Mutant data in the IMPC has a special structure, resulting from mice being born in the same litters and being phenotyped closely together in time (batch effect), which must be replicated in the resampling approach. We address this by utilising *structured resampling* that replaces the mutants with the closest random controls in time. We create artificial mutant groups by randomly sliding the true mutant structure over the time domain of controls, collecting as many controls as there were mutants in the original set, and repeating this procedure five times per dataset (supplemental Figure 1 shows an illustration of three iterations of the structured resampling on the *Bone Mineral Content* parameter).

The outcome of the simulation study consists of 18 IMPC procedures across 11 centres and over 2.5 *million* analyses and p-values. Comparing the results from the IMPC standard and soft windowed analyses on resampled data, we detect an overall of 14,201 and 12,716 false positives (FP), respectively, at the signficance level used by the IMPC, 0.0001. This constitutes more than a 10% relative improvement in FPs when the soft windowed method is applied. Table 1 shows the top ten IMPC procedures with the significant changes in the FPs. From this table, the procedures *Body Composition*, *Open Field*, *Urinalysis*, *Heart Weight*, *Acoustic Startle and Pre-pulse Inhibition* account for the highest relative reduction of 68% in FPs, whereas the *Clinical Blood Chemistry*, *X-Ray*, *Insulin Blood Levels*, *Electrocardiogram* and *Eye Morphology* account for the maximum increase of 32% in FPs. Supplemental Figure 2 shows parameters from the Body Composition and Clinical Blood Chemistry procudures that showed the biggest loss and gain in false positives for assocaited data parameters, respectively. This plot shows an improvement in decreasing FPs in all IMPC_DXA parameters, which contrasts with an increase in the FPs for IMPC CBC parameters. We further examined the top two IMPC_CBC parameters, *Alanine aminotransferase* (IMPC_CBC_013) and *Aspartate aminotransferase* (IMPC_CBC_012) in Supplemental Figure 3, and noted a high level of randomly deviated points from the mean of controls that can bias the outcome of the structured resampling.

**Table 1:** Top ten IMPC procedures with the highest change in the total number of false positives

| *Procedure name* | *Procedure id* | *Total p-values* | *NFP*<sup>*</sup> | *WFP*<sup>†</sup> | *WFP/(NFP+WFP)%* |
| --- | --- | --- | --- | --- | --- |
| Body Composition (DEXA lean/fat) | IMPC_DXA | 167789 | 3809 | 2293 | 37.58 |
| Clinical Blood Chemistry | IMPC_CBC | 320949 | 1472 | 2414 | 62.12 |
| Open Field | IMPC_OFD | 182894 | 1507 | 830 | 35.52 |
| Haematology | IMPC_HEM | 243640 | 3125 | 2746 | 46.77 |
| Heart Weight | IMPC_HWT | 16236 | 553 | 409 | 42.52 |
| Acoustic Startle and Pre-pulse Inhibition (PPI) | IMPC_ACS | 73177 | 352 | 243 | 40.84 |
| X-ray | IMPC_XRY | 7016 | 27 | 135 | 83.33 |
| Insulin Blood Level | IMPC_INS | 9465 | 63 | 164 | 72.25 |
| Electrocardiogram (ECG) | IMPC_ECG | 122257 | 378 | 471 | 55.48 |
| Eye Morphology | IMPC_EYE | 15739 | 86 | 153 | 64.02 |

\* False positives from the non-windowed results
† False positives from the soft-windowed results

### Soft windowing as part of the IMPC statistics pipeline

We next show the performance of the soft-windowing approach on IMPC data by integrating it into the standard IMPC statistics pipeline, PhenStat ^34^. Using data release 9.2 (January 2019), we re-analysed 14 *million* + data points from which 10 *million* + are mutant animals across the range of IMPC phenotyping procedures. The original IMPC standard analysis that did not apply the soft windowing approach to select the control data encompassed 403,000+ analyses and p-values. This analysis led to 12,728 significant p-values (< 0.0001), compared to 16,415 significant p-values when the soft windowing was applied, an increase of 30% in total significant p-values. The IMPC assigns mouse lines with phenotype terms from the Mouse Phenotype Ontology when a significant deviation from the control data is detected for a given data parameter ^35^. Our windowing approach led to 17,391 associations gained and 15,996 associations lost. To explore these differences further, we created an online tool that displays the entire control dataset for a given mouse line-parameter assay with the statistical summaries for both the non-windowed methodology and the soft windowed approach. Users may filter on a number of attributes, arrange filter order, zoom in on data visualisation, or navigate directly to the results (https://wwwdev.ebi.ac.uk/mi/impc/dev/phenotype-archive/media/images/windowing/).

Figure 5 shows the corresponding visualization on the IMPC website of the data previously shown in Figure 1 (left) for the *Forelimb grip strength normalised against body weight* parameter from the IMPC Grip Strength procedure. The soft window is indicated, as well as changes in the total number of controls (here 1,572 fewer after soft windowing). Further, the p-value corresponding to the genotype effect shows a significant change in magnitude, from 2.05 × 10^−4^ to 6.75 × 10^−18^ after applying the soft windowing.

**Figure 5.**
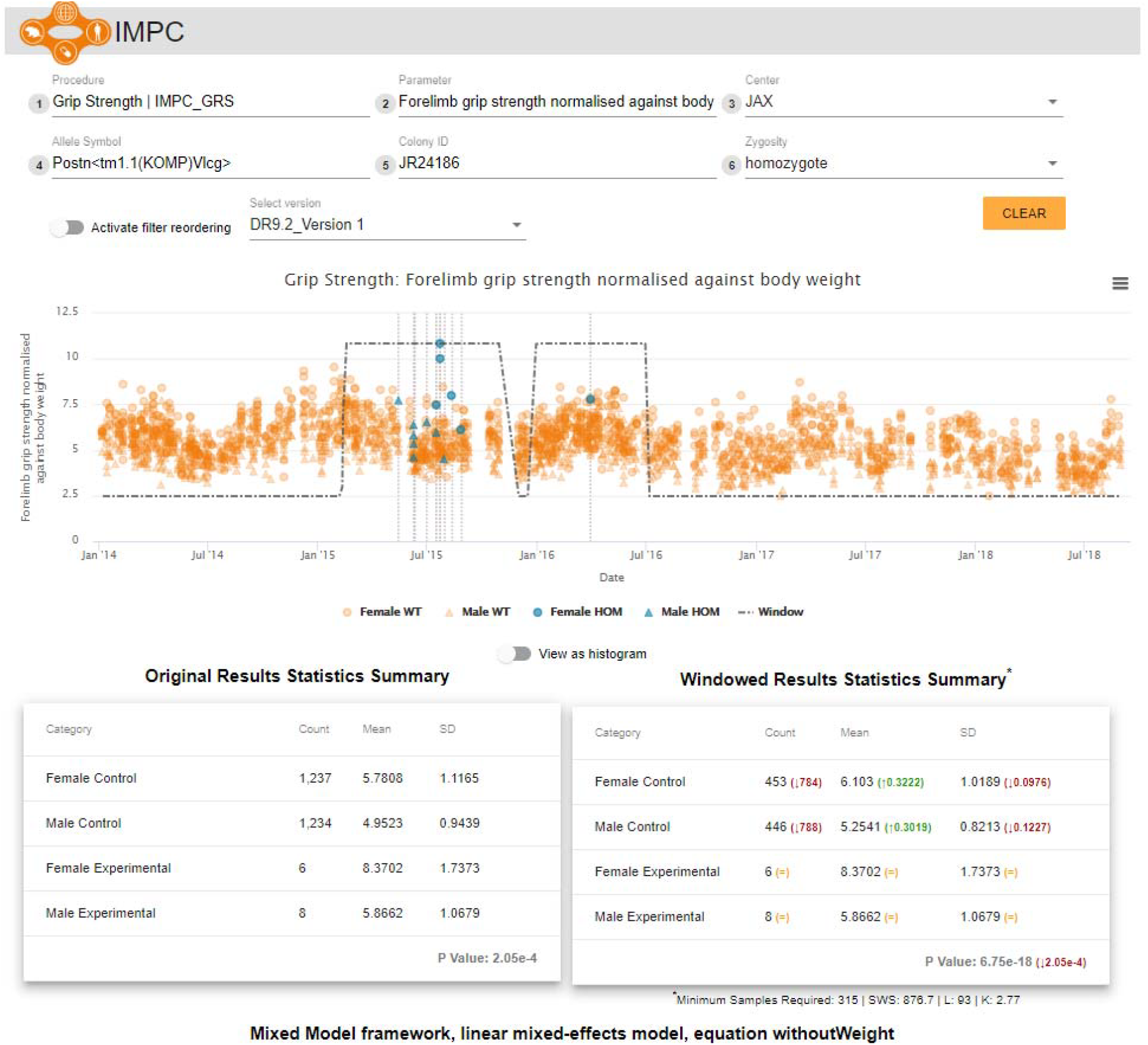
The soft windowing visualization in the IMPC website for the *Forelimb grip strength normalised against body weight* from the IMPC *Grip Strength* procedure. The plot shows the response over time as well as the fitted soft windows. The tables below show the comparison between the descriptive statistics obtained from the standard (non-windowed) analysis on the left and the soft windowed approach on the right. The p-values correspond to the genotype effect after applying the statistical analyses taking the corresponding controls based on the non-window and soft windowed approaches, respectively.

We then tested if our soft windowed analysis changed our human disease model discovery rate. We have previously described the IMPC Phenodigm translational pipeline that automatically detects phenotypic similarities between the IMPC strains and over 7,000 rare diseases described in the Online Mendelian Inheritance in Man (OMIM), Orphanet and the Deciphering Developmental Disorders (DDD) databases ^35^. This pipeline generates qualitative scores on how well a mouse line’s associated phenotypes overlap with the phenotypes of the human rare disease populations ^35–40^. By comparing the disease model resulting from our soft windowed analysis vs non-windowed analysis for IMPC data release 9.2, we find a slight increase in the number of disease models (106 vs 99 models using a threshold of 50% phenotype overlap from a set of 2,082 mouse lines that contain mutations-Supplemental Table I).

## Discussion

High-throughput phenomics is a powerful tool for the discovery of new genotype-phenotype associations and there is an increasing need for innovative analyses that make effective use of the voluminous data being generated. Batch effects are inevitable when a large amount of data is collected at different times and/or sites and, therefore, need to be accounted for in the statistical analysis. In this study, we developed a novel “soft windowing” method which selects a window of time to include controls that are locally selected with respect to experimental animals, thus reducing the noise level in the data collected over long periods of time (years). Soft windowing has notable advantages over a more traditional hard windowing approach. In contrast to the limited data points included in the hard windowing method, the entire dataset is considered for the analysis. To this end, we engineered a weighting function to produce weights in the form of a window of time. Control data collected proximally to mutants were assigned the maximal weight, while data collected earlier or later had less weight. This method has the capability of producing indivdual windows as well as merging intersected ones. Moreover, the method was implemented to automatically select window size and shape.

The performance of the method was shown on a simulated scenario that uses real control data collected by the IMPC high-throughput pipelines to assess detection of false positives. We also showed the enhancements to the IMPC statistical pipeline that establishes genotype-phenotype associations by comparing mutants vs control data using our soft windowed approach.

There are two known conditions that affect the method: (1) The weight generating function can be slow when there are too many (> 20) distinct windows, however, we have optimised the algorithm to be fast enough for the typical IMPC number of peaks (≈3 seconds for 1500 samples and 16 peaks under *k* = 1 and *l* = 30); and (2) Our resampling scenario indiciated that our soft windowing approach is sensitive to the data that has a high level of outliers or random deviation from the mean. This may result from a bias in the design of the resampling but may also indicate that using all available controls maybe appropriate for the cases with extreme variability.

Our soft windowing approach addresses the scaling issues associated with analysing an ever-increasing set of control data in long-term projects by eliminating controls with weights sufficiently close to zero from future analysis. In the case of the IMPC, once a window of control data is determined for a dataset, there would be no further requirement to re-analyse the dataset with each subsequent data release. This will reduce the computational resources needed with the resulting gene-phenotype associations remaining stable, greatly facilitating data exchange with research groups trying to functionally validate genes and their disease variants. Our findings also have important implications for such efforts as the UK BioBank and the All of Us initiatives where large cohort sizes coupled with mobile medical sensors are generating phenotype data at an unprecedented rate ^41,42^. Researchers performing restrospective analysis to analyse exposures for a defined outcome group (e.g. metabolic disease) are challenged by the variability and longitudinal characteristics associated with these datasets. The methods described here can be used with these human health resources to maximise analytical power and help researchers find the genetic and environmental contributers to human diseases.

