## Supplementary materials for "Soft Windowing Application to Improve Analysis of High-throughput Phenotyping Data"

Supplementary figure 1: The scatter plot of the data from the *Bone Mineral Content* parameter from the IMPC *Body Composition* (IMPC DXA) procedure for a selected mutant line and its corresponding controls. The top plot shows the true structure of the mutants that is used to construct the structured resampling. The bottom plot shows three iterations of the structured resampling.

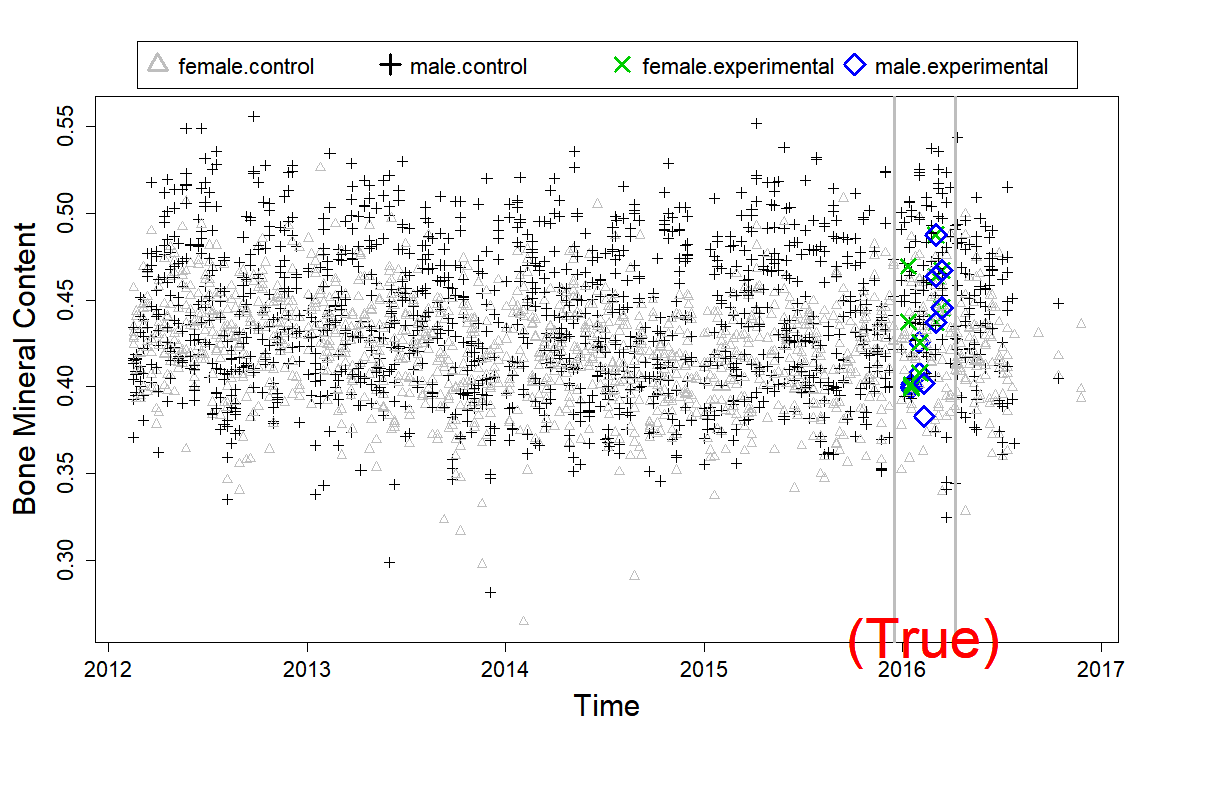

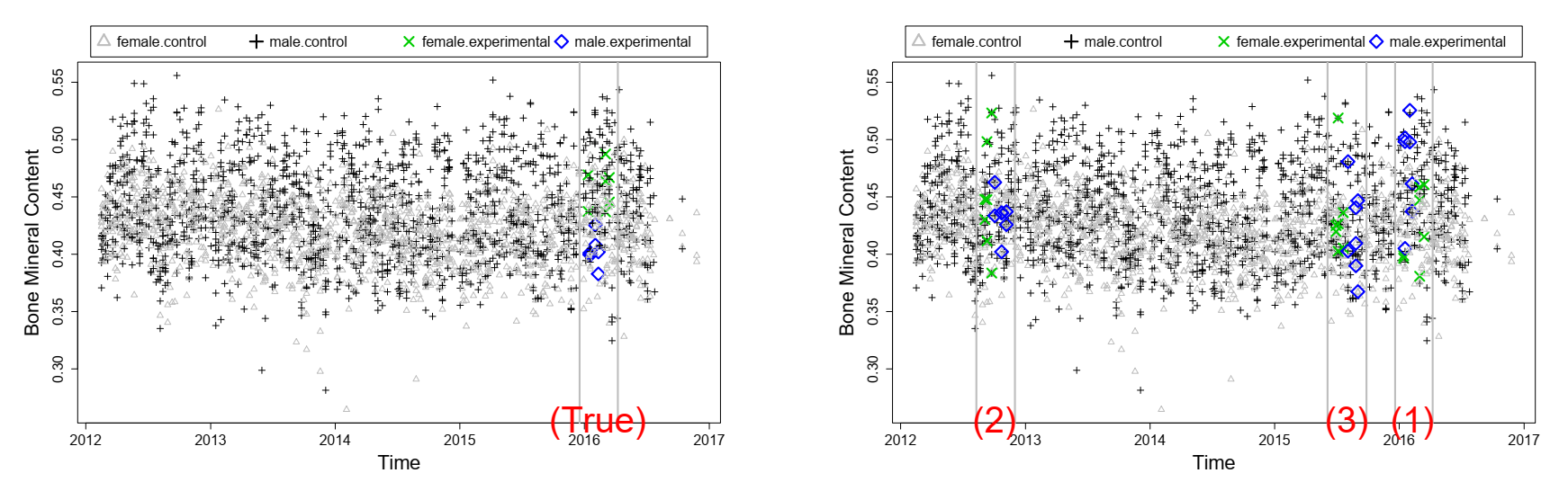

Supplementary figure 2: Number of false positives detected in the phenotypic parameters measured as part of the IMPC DXA (body composition) and IMPC CBC (clinical blood chemistry) procedures when re-sampling controls and applying the non-windowed analysis (black) and the soft-windowed analysis (red).

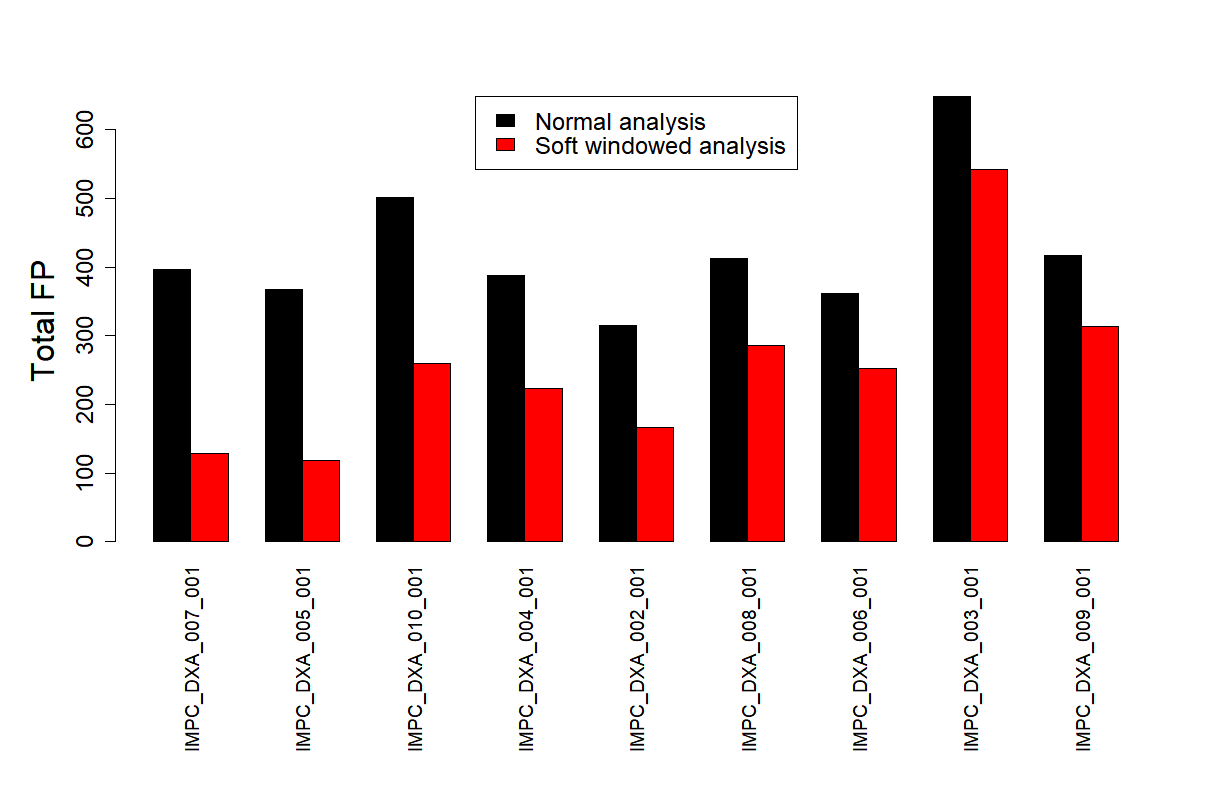

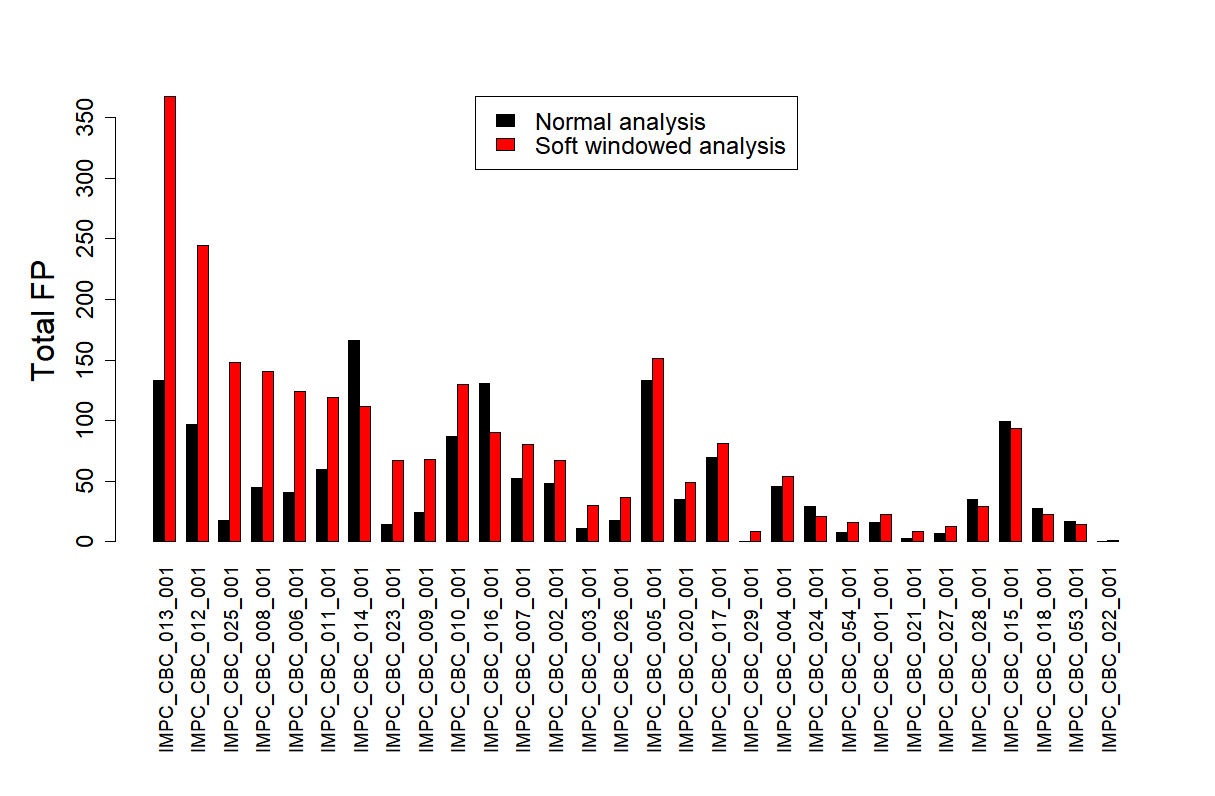

Supplementary figure 3: The plots show the deviation from the mean and variability of the controls over time for two parameters, namely *Alanine aminotransferase* (IMPC_CBC_013) and *Aspartate aminotransferase* (IMPC_CBC_012), that are parts of the IMPC *Clinical Blood Chemistry* (IMPC_CBC) procedure.

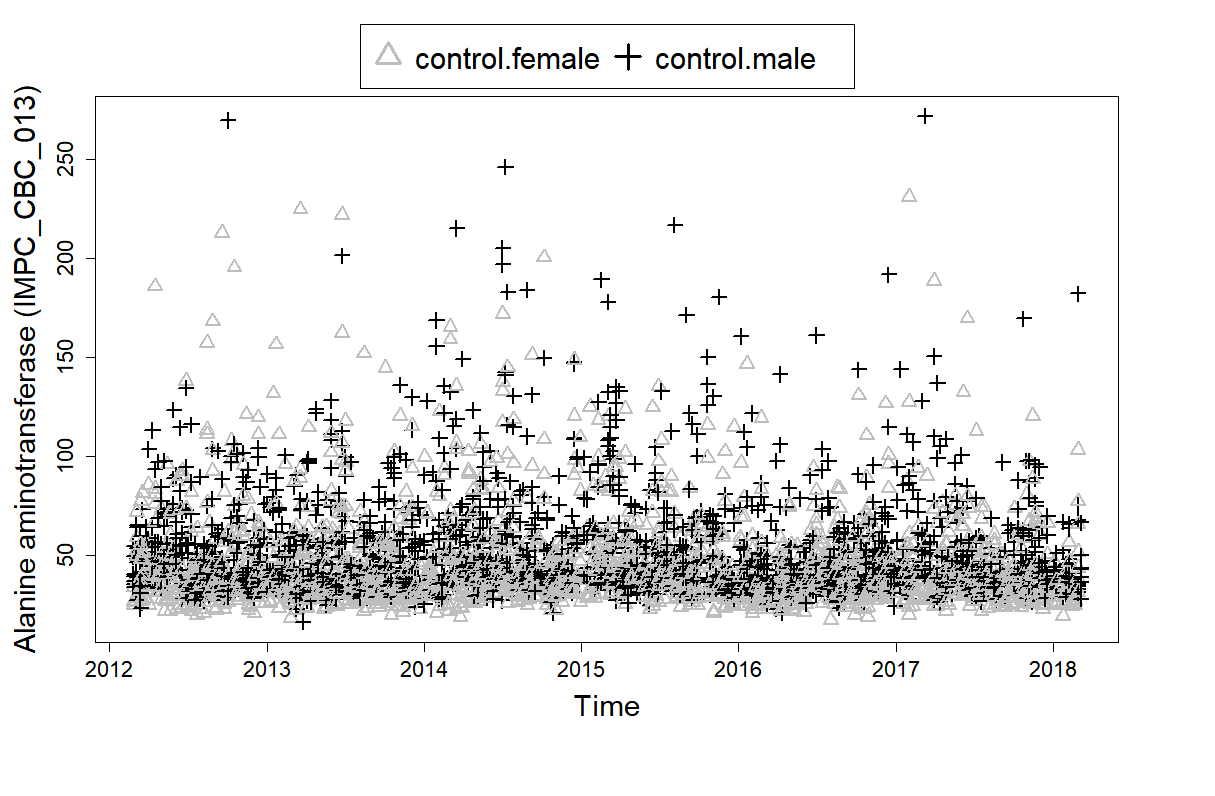

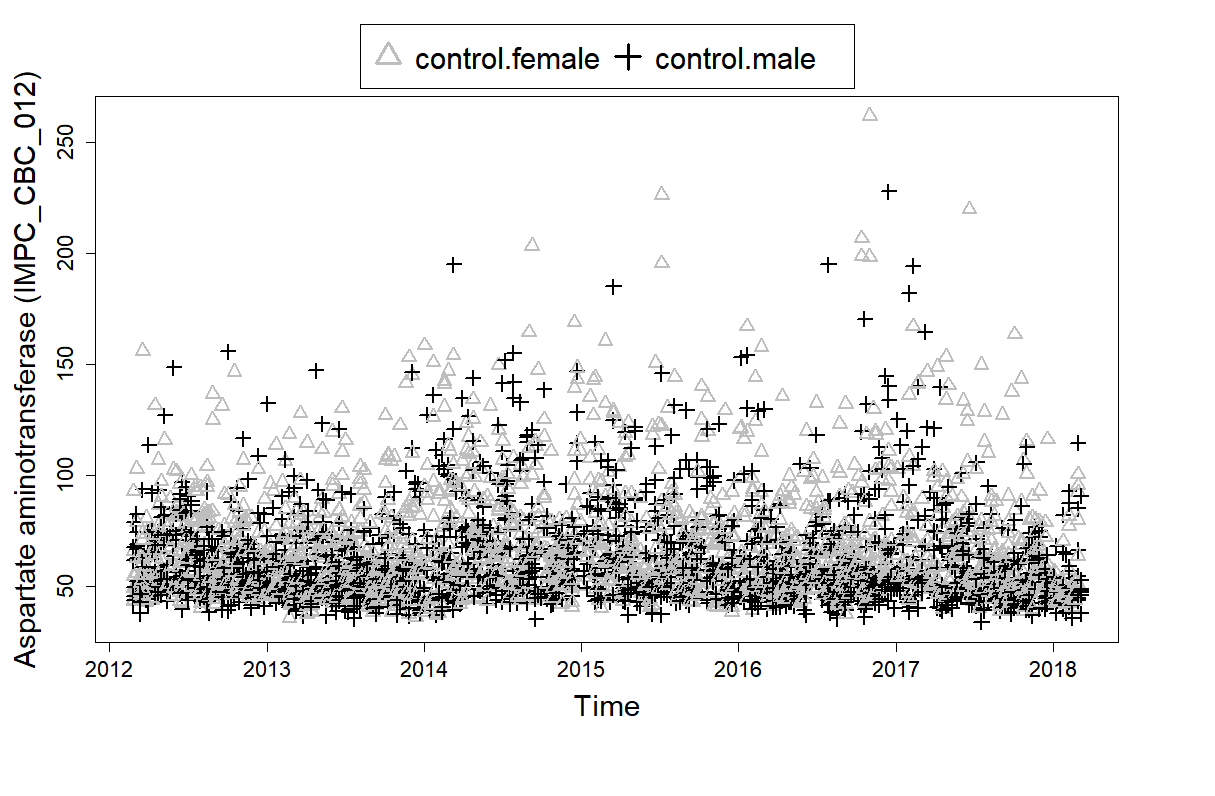
